# Distinct hippocampal mechanisms support concept formation and updating

**DOI:** 10.1101/2024.02.14.580181

**Authors:** Michael L. Mack, Bradley C. Love, Alison R. Preston

## Abstract

Learning systems must constantly decide whether to create new representations or update existing ones. For example, a child learning that a bat is a mammal and not a bird would be best served by creating a new representation, whereas updating may be best when encountering a second similar bat. Characterizing the neural dynamics that underlie these complementary memory operations requires identifying the exact moments when each operation occurs. We address this challenge by interrogating fMRI brain activation with a computational learning model that predicts trial-by-trial when memories are created versus updated. We found distinct neural engagement in anterior hippocampus and ventral striatum for model-predicted memory create and update events during early learning. Notably, the degree of this effect in hippocampus, but not ventral striatum, significantly related to learning outcome. Hippocampus additionally showed distinct patterns of functional coactivation with ventromedial prefrontal cortex and angular gyrus during memory creation and premotor cortex during memory updating. These findings suggest that complementary memory functions, as formalized in computational learning models, underlie the rapid formation of novel conceptual knowledge, with the hippocampus and its interactions with frontoparietal circuits playing a crucial role in successful learning.

**Significance statement:** How do we reconcile new experiences with existing knowledge? Prominent theories suggest that novel information is either captured by creating new memories or leveraged to update existing memories, yet empirical support of how these distinct memory operations unfold during learning is limited. Here, we combine computational modeling of human learning behaviour with functional neuroimaging to identify moments of memory formation and updating and characterize their neural signatures. We find that both hippocampus and ventral striatum are distinctly engaged when memories are created versus updated; however, it is only hippocampus activation that is associated with learning outcomes. Our findings motivate a key theoretical refinement that positions hippocampus is a key player in building organized memories from the earliest moments of learning.

## Introduction

Learning often relies on integrating new experiences with existing knowledge to encode regularities. Yet not all experiences align with what we know. A child learning about bats will discover that, despite having wings, they are mammals, not birds. Classic learning theories propose that such experiences are learned like any other: abstracted into a prototype (Homa et al., 1973) or encoded as exemplars (Nosofsky, 1986). Alternatively, distinct memory operations may be triggered throughout learning depending on how new information matches prior experiences (Love et al., 2004; Mack et al., 2018; Morton and Preston, 2021). Existing memories may be updated to generalize across related experiences, or new memories may be created to distinctly capture novel ones. Such learning may support precise conceptual discrimination (e.g., the child can preserve general knowledge about mammals while distinctly encoding bats) allowing for more flexible future inferences. However, identifying when these qualitatively different memory operations occur during learning has proven empirically challenging. Here, we combine computational modeling with human neuroimaging to quantify, moment-by-moment, the distinct memory mechanisms that support successful concept learning.

Seminal findings identified multiple systems underlying learning (Knowlton et al., 1996; Poldrack et al., 2001; Shohamy et al., 2008). Hippocampal engagement was thought to capture episodic details early in learning, while basal ganglia incrementally learned stimulus-response associations that reflected behaviour. This view has evolved to emphasize the hippocampus’s flexibility in building structured memories (McClelland et al., 1995; O’Reilly and Rudy, 2001; Schapiro et al., 2017; Duncan and Schlichting, 2018; Mack et al., 2018; Sučević and Schapiro, 2023). Specifically, hippocampal comparator processes evaluate overlap between new experiences and existing knowledge to trigger either memory integration—updating stored representations with new information (Schlichting et al., 2015)—or memory differentiation—creating distinct representations for conceptually novel experiences (Hulbert and Norman, 2014; Ritvo et al., 2023). The transient nature of these hippocampal memory operations is evident in animal models: moment-by-moment fluctuations in slow and fast gamma oscillations between hippocampal subfields and entorhinal cortex (Colgin et al., 2009; Bieri et al., 2014; Igarashi et al., 2014) and distinct sharp-wave ripples in hippocampus (Singer et al., 2013; Joo and Frank, 2018) reflect behaviourally relevant state changes in memory function.

In humans, evidence for such hippocampal state changes is emerging. Mismatch between cued memories and current experience increases hippocampal activation (Duncan et al., 2012) and alters connectivity patterns (Bein et al., 2020), and hippocampal representations of associative pairs become markedly differentiated once learned (Wanjia et al., 2021). Trial-by-trial shifts in decision certainty during category learning (Davis et al., 2012a; Bowman and Zeithamova, 2018; Theves et al., 2021) are also linked to hippocampal activation. The key open question is whether distinct memory creation and updating operations, as they unfold throughout learning, are directly reflected in hippocampal function.

We address this gap with SUSTAIN (Love et al., 2004), a computational learning model that explicitly predicts *when* and *how* new information is encoded into memory. SUSTAIN’s mechanisms for representation learning parallel theorized hippocampal operations (Love and Gureckis, 2007; Davis et al., 2012a; Mack et al., 2016, 2018). The model learns via adaptive clustering: clusters (i.e., memory traces) of stimulus features are created or updated depending on the match between existing memories—constructed from prior experiences and response history—and the current stimulus. SUSTAIN thus provide a principled means to identify when specific memory operations occur throughout learning to build flexible memory representations.

Here, we tested the prediction that brain regions involved in adaptive memory are distinctly engaged during SUSTAIN-predicted memory updating and creation. We recorded functional magnetic resonance imaging (fMRI) data while participants completed four visual category learning tasks, derived model-based trial-by-trial memory operation predictions, and interrogated brain activation through the lens of these model predictions. We demonstrate that hippocampus and ventral striatum reflect dynamic memory operations during early learning, but only hippocampal activation predicts learning outcomes.

## Materials and Methods

### Participants

Twenty-five volunteers (13 females, mean age 21.6 years old, ranging from 18 to 29 years) participated in the experiment. All subjects were right-handed, had normal or corrected-to-normal vision, and were compensated $75 for participating.

### Stimuli

Four stimulus image sets were used in the experiment (**Figure 1A**). Each stimulus set included eight images consisting of all combinations of three binary stimulus features (flower: outer petal shape, inner petal shape, centre colour; fribbles: tail, ears, legs; amoeba: cell structure features in the 3 cell arms; spaceships: wings, body, antennae).

**Figure 1.**
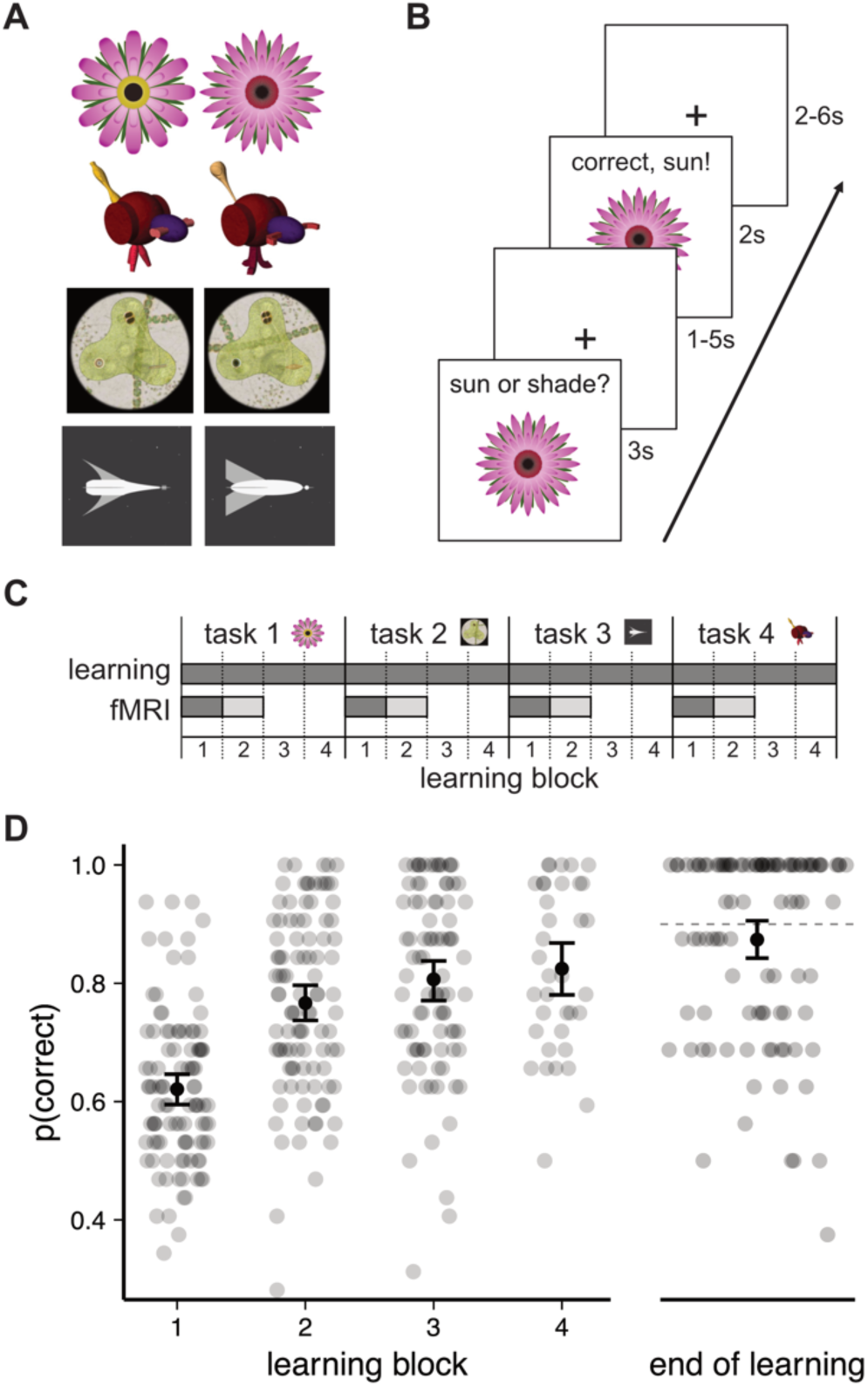
Learning tasks stimuli and trial schematic and learning performance. **A)** The four stimulus sets consisted of visual objects composed of three binary feature dimensions (Table 1). **B)** Learning trials followed typical feedback-based learning paradigm: a stimulus was shown for 3s during which participants could make a categorization response. After a variable delay (1-5s), feedback including the stimulus, whether or not the response was correct, and the correct category was presented for 2s followed by a variable delay (2-6s) before the next trial. **C)** Experimental sessions consisted of four learning tasks each with four blocks of feed-back learning trials. fMRI data was collected during the first two learning blocks. Analyses were restricted to blocks with darker shading (i.e., only block 1 of fMRI data). **D)** The probability of a correct response is plotted separately for each learning block and end of learning. Group averages and bootstrapped 95% confidence intervals are plotted in black; accuracy for individual participants in each of the four tasks are plotted in the smaller transparent points. A learning task would end after blocks 2 or 3 if the participant reached threshold of 90% correct on last 16 trials; this is reflected in fewer points plotted in blocks 3 and 4. End of learning accuracy (right) is depicted for each participant separately for each task and includes performance for tasks in which participants reached threshold in earlier learning blocks. The dotted line depicts the 90% learning threshold. Data includes N=25 participants each in 4 tasks.

**Table 1:** Stimulus features and class associations for the three learning problems. Each of the eight stimuli are represented by the binary values of the three feature attributes. The stimuli are assigned to different classes (A or B) across the three category structures according to rules that depend on a combination of feature attributes.

| stimulus | feature attribute |  |  | category structure |  |  |
| --- | --- | --- | --- | --- | --- | --- |
|  | 1 | 2 | 3 | type 3 | type 4 | type 5 |
| 1 | 0 | 0 | 0 | B | B | B |
| 2 | 0 | 0 | 1 | B | B | B |
| 3 | 0 | 1 | 0 | B | B | B |
| 4 | 0 | 1 | 1 | A | A | A |
| 5 | 1 | 0 | 0 | A | B | A |
| 6 | 1 | 0 | 1 | B | A | A |
| 7 | 1 | 1 | 0 | A | A | A |
| 8 | 1 | 1 | 1 | A | A | B |

There were two versions of each (e.g., flower had pointy or round outer petals). Following procedures from previous work using the amoeba stimuli (Blair et al., 2009), the eight amoeba stimuli were randomly presented on different backgrounds throughout learning. All feature versions are depicted in Figure 1A. All stimulus images used in the experiment are available on OSF (https://osf.io/jk79v/).

### Procedure for learning tasks

After an initial screening and consent in accordance with the University of Texas Institutional Review Board, participants were instructed on the category learning problems. Participants then performed the problems in the MRI scanner by viewing visual stimuli back projected onto a screen through a mirror attached onto the head coil. Foam pads were used to minimize head motion. Stimulus presentation and timing was performed using custom scripts written in Matlab (Mathworks) and Psychtoolbox (www.psychtoolbox.org) on an Apple Mac Pro computer running OS X 10.7.

Participants were instructed to learn to classify the stimuli based on the combination of the features using the feedback displayed on each trial. As part of the initial instructions, participants were made aware of the three features and the two different values of each feature across all four stimulus sets. Before beginning each learning task, additional instructions that described the cover story for the current task and which buttons to press for the two stimulus categories were presented to the participants. One example of this instruction text is as follows: “Each flower prefers either Sun or Shade to grow best. The environment that each flower prefers depends on one or more of its features. On each trial, you will be shown a flower and you will make a response as to that flower’s preferred environment. Press the ‘1’ button under your index finger for Sun or the ‘2’ button under your middle finger for Shade.” After the instruction screen, the two fMRI scanning runs (described below) for that problem commenced, with no further problem instructions. After the two scanning runs for a problem finished, the participants continued the learning task until criterion was reached or all trials for a task were completed. The next learning task began with the corresponding cover story description. Importantly, the rules that defined the learnings tasks were not included in any of the instructions; rather, participants had to learn these rules through trial and error.

The category structures underlying the learning tasks correspond with Shepard et al.‘s problem types 3, 4, and 5 (Shepard et al., 1961) (**Figure 1A**, **Table 1**). These structures were selected because optimally learning them requires attending to all three feature dimensions and both generalizing across and distinctly representing specific stimuli (Shepard et al., 1961; Love et al., 2004). In other words, memory creation and updating are key processes for learning these tasks. Notably, simulations of behaviour in these tasks with SUSTAIN suggest that memory creation operations are relatively infrequent and tend to occur early in learning; as such we included four variants of the learning task to capture enough of these create trials with a balanced number of update trials.

The binary values of the feature attributes along with the class association for the three category structures are depicted in Table 1. The stimulus features were randomly mapped onto the attributes for each participant and task. For each participant, the four stimulus sets were randomly paired with the three category structures with one structure repeated. The order of the tasks was randomly shuffled across participants.

The learning tasks consisted of learning trials (**Figure 1B**) during which a stimulus image was presented for 3s. During stimulus presentation, participants were instructed to respond to the stimulus’ category by pressing one of two buttons on an fMRI-compatible button box. Stimulus images subtended 7.3° × 7.3° of visual space. The stimulus presentation period was followed by a 1-5s fixation. A feedback screen consisting of the stimulus image, text of whether the response was correct or incorrect, and the correct category was shown for 2s followed by a 2-6s fixation. The timing of the stimulus and feedback phases of the learning trials was jittered to optimize general linear modeling estimation of the fMRI data. Within one functional run, each of the eight stimulus images for a given task was presented in four learning trials. The order of the learning trials was pseudo randomized in blocks of 16 trials such that the eight stimuli were each presented twice. Across a learning task, each stimulus was presented 8-16 times depending on whether or not a participant reached learning criterion (see below). One functional run was 388s in duration. In total, participants completed four learning tasks (**Figure 1C**). Each of the learning tasks included two functional runs corresponding to the first 64 trials of the task. After these two functional runs, learning trials continued for two more blocks of 32 trials without fMRI scanning and with no fixation periods after stimulus presentation or feedback. If participants reached 90% accuracy on the final 16 trials of learning blocks 2, 3, or 4, the task ended. Otherwise, participants completed 128 trials for a task. The entire experiment lasted approximately 75 minutes.

### Learning performance

Performance during the learning tasks was characterized within each learning block as the average accuracy across the block’s 32 trials (**Figure 1D**). During data collection, if a participant reached or surpassed 90% accuracy on the final 16 trials of blocks 2, 3, or 4, that task was ended. That is, some participants learned quickly and finished tasks early. As such, end of learning performance was characterized by average accuracy on the final 16 trials that the participants completed (**Figure 1D**). Across all tasks and participants, the average end of learning accuracy was 87.4% (median: 93.75%, SD: 15.8%, range: 37.5-100%) with a majority reaching the 90% threshold (57/100 completed tasks). Given that end of learning performance was highly skewed, we classified task performance by whether or not the 90% accuracy threshold was reached. This “learner” label was subsequently leveraged in relating neural indices of memory formation to end of learning performance.

### Computational learning model

Participant behavior was modeled with an established mathematical learning model, SUSTAIN (Love et al., 2004). SUSTAIN accounts for behaviour in category learning tasks with an adaptive clustering mechanism (**Figure 2A**) in which clusters (i.e., memory traces) of stimulus features and their association to a category label are created or updated depending on the match between the model’s existing memories and the current stimulus information. Generalization occurs through incremental updates of existing clusters, whereas differentiation arises via cluster creation. Notably, SUSTAIN’s adaptive clustering mechanism allows for multiple clusters to be linked to the same category label, thereby endowing it with a distinct ability to learn complex category structures.

**Figure 2.**
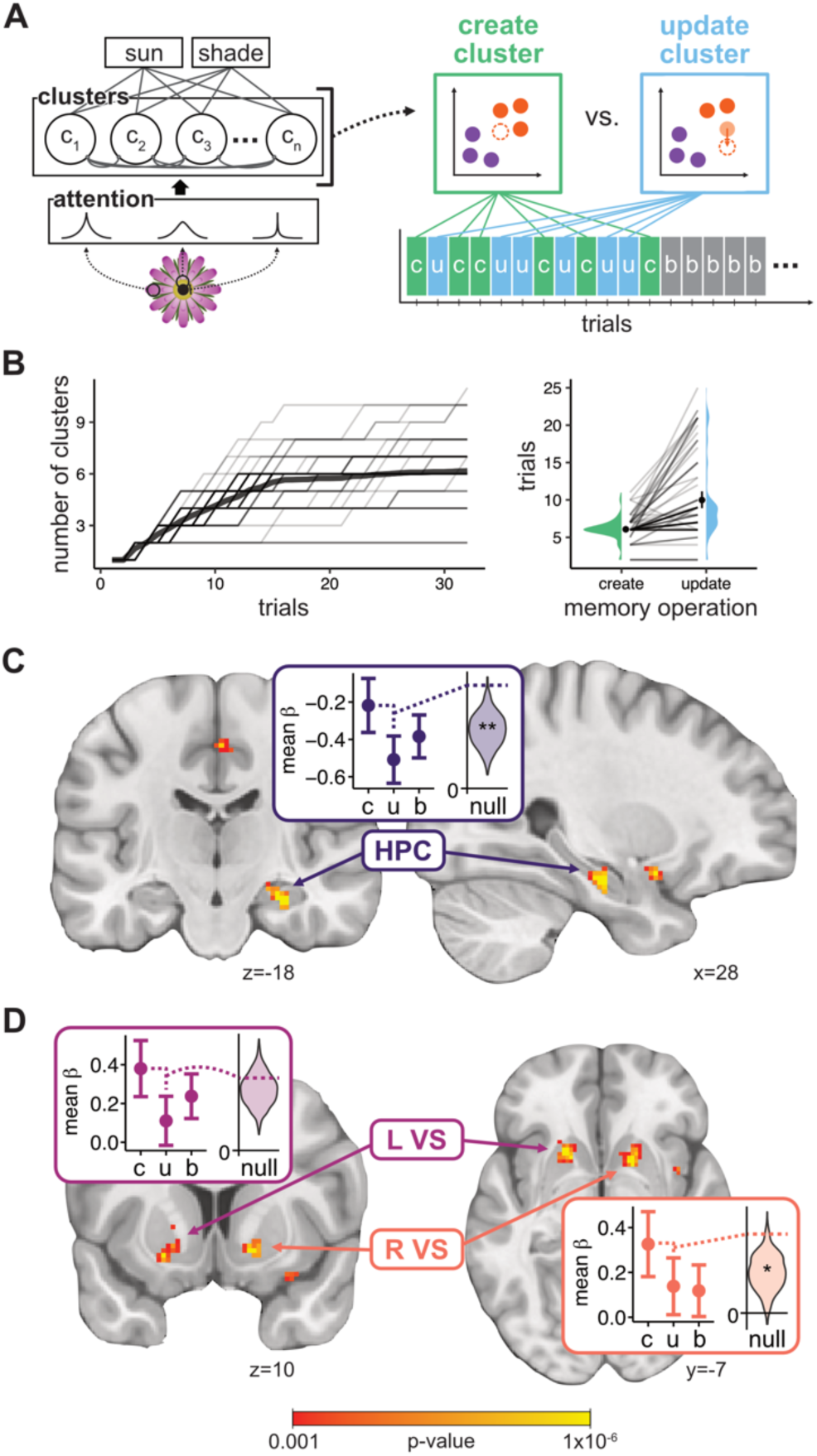
Illustration of the SUSTAIN computational model and model-based fMRI analyses of memory formation functions. **A)** SUSTAIN formalizes learning as the interaction between feature-based selective attention and memory representations formed via a clustering mechanism. On each learning trial a new cluster is created (green *c*) or an existing cluster is updated (blue *u*) depending on the match of the current stimulus to existing clusters. As learning progresses clusters become effectively fixed (grey *b*). **B)** Model simulations of learning behaviour (left plot) showed that clusters were created throughout the initial trials (participant-specific traces shown in transparent thin lines, average trace depicted as solid black line). Average number of create and update trials per participant in each task is depicted on the right with. Error bars represent 95% CI and lines connect points from individual participants in the different tasks. **C)** A region in right hippocampus (HPC, peak [30, -18, -19]) showed higher activation for create relative to update trials. Inset plot depicts mean β estimates at region peak for create (c), update (u), and baseline (b) trials. Error bars represent 95% confidence intervals. Violin plots depict shuffled null distribution of create-update effect (*z* statistic) with observed effect. **D)** Two regions showing greater create than update engagement were localized to left and right ventral striatum (L VS, peak [-20, 14, -8]; R VS, peak [8, 20, -4]). Clusters were defined by a voxel wise threshold of p=0.001 and cluster-extent threshold of p=0.05 (>42 voxels). N=25. **p<0.01, *p<0.05

Stimuli are encoded by SUSTAIN into perceptual representations based on the value of the stimulus features. The values of these features are biased according to attention weights operationalized as receptive fields on each feature attribute. During learning, these attention weight receptive fields, which change as a function of the latent model variable *1_i_*, are tuned to give more weight to diagnostic features. SUSTAIN represents knowledge as memory clusters of stimulus features and class associations that are built and tuned over the course of learning. Key to the current work is SUSTAIN’s mechanism for modifying these clusters throughout learning. Specifically, on each trial, attention-weighted feature values for the current stimulus are compared to existing clusters. If a cluster matches the stimulus well, that cluster drives the decision process and is subsequently *updated* to reflect the attention-weighted feature values of the current stimuli. If there is a poor match between the stimulus and the existing clusters, a new cluster is *created* that reflects the feature values of the current stimulus. Throughout learning, therefore, new clusters are created, and existing clusters updated depending on the trial sequence and category structure of the current learning task. A full mathematical formulization of SUSTAIN is provided below:

### Perceptual encoding

An input stimulus is presented to SUSTAIN as a pattern of activation on input units that code for the different stimulus features and possible values that these features can take. For each stimulus feature, *i* (e.g., a flower’s outer petals), with *k* possible values (e.g., pointy or rounded petals), there are *k* input units. Input units are set to one if the unit represents the feature value or zero otherwise. The entire stimulus is represented by *I ^posik^*, with *i* indicating the stimulus feature and *k* indicating the value for feature *i*. “pos” indicates that the stimulus is represented as a point in a multidimensional space. The distance *μ_ij_* between the *i*th stimulus feature and cluster *j*’s position along the *i*th feature is

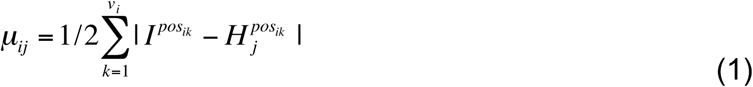

such that *v_i_* is the number of possible values that the *i*th stimulus feature can take and is cluster *j*’s position on the *i*th feature for value *k*. Distance *μ_ij_* is always between 0 and 1, inclusive.

### Response selection

After perceptual encoding, each cluster is activated based on the similarity of the cluster to the input stimulus. Cluster activation is given by:

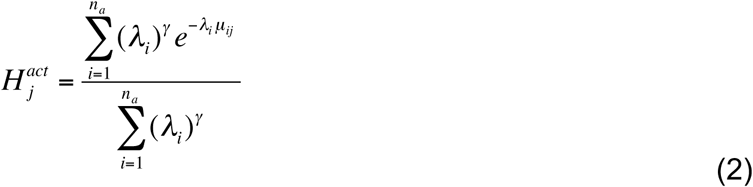

where 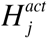 is cluster *j*’s activation, *n*_a_ is the number of stimulus features, *λ_i_* is the attention weight receptive field tuning for feature *i*, and γ is the attentional parameter (constrained to be non-negative). Clusters compete to respond to an input stimulus through mutual inhibition. The final output of each cluster *j* is given by:

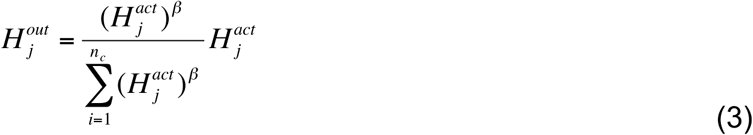

where *n_c_* is the current number of clusters and β is a lateral inhibition parameter (constrained to be non-negative) that controls the level of cluster competition. The cluster that wins the competition, *H_m_*, passes its output to the *k* output units of the unknown feature dimension *z*:

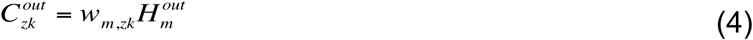

where 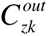is the output of the unit representing the *k*th feature value of the *z*th feature, and w*_m,zk_* is the weight from the winning cluster, *H_m_*, to the output unit *C_zk_*. In the current simulations, the class label is the only unknown feature dimension. Thus, equation 4 is calculated for each of the two values of the class label. Finally, the probability of making a response *k* for a queried dimension, *z*, on a given trial is:

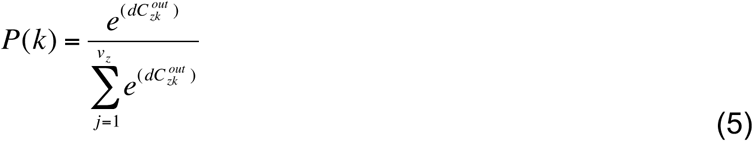

Note that a response scaling parameter *d* is included in equation 5 to model probabilistic versus deterministic choices; when *d* is high, accuracy is stressed and the cluster with the largest output is almost always chosen.

### Memory cluster modification

SUSTAIN was initialized with zero clusters. During learning, clusters are recruited in response to a combination of the order of the stimuli presented in the participant-specific trial orders and the error feedback received on each trial. Two events could lead to the creation of a new cluster: 1) the model predicts the incorrect class label or 2) the winning cluster’s activation is below a threshold, τ (constrained to be between 0 and 1). If either of these two criteria are true, a new cluster is created; otherwise, the winning cluster from the cluster competition is updated to reflect current stimulus features and class label according to the learning rules explained next.

### Learning

SUSTAIN’s learning rules determine how clusters are updated during learning. Only the winning clusters are updated. If a new cluster is recruited on a trial, it is considered the winning cluster. Otherwise, the cluster that is most similar to the current stimulus will be the winner. The winning cluster *H_m_*, is adjusted by:

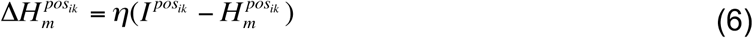

where ρι is the learning rate parameter. The result of the updating is that the winning cluster moves toward the current stimulus. Over the course of learning, each cluster will tend toward the center of its members. Attention weight receptive field tunings for the different feature dimensions are updated according to:

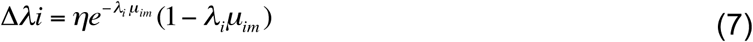

where *m* indexes the winning cluster. The weights from the winning cluster to the output units are adjusted by a one-layer delta learning rule.

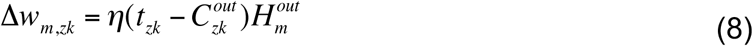

### Simulations

Stimuli were presented to SUSTAIN using the same trial order for each task as experienced by the participants. Before each task, the attention weight receptive field tunings and clusters were reinitialized. For each participant and task, the free parameters, *γ*, β, η, *d*, and *τ*, were optimized to best match the participant’s trial-by-trial responses. Specifically, SUSTAIN’s predicted probability of a making the same response as the participant (**Eq. 5**) was summarized with log likelihood (*lnL*) and the model parameters were optimized to maximise likelihood using a differential evolution genetic algorithm approach (Storn and Price, 1997; Mack et al., 2016) (*scipy* version 1.2.1). To ensure that these trial-by-trial model predictions were successful, we compared the resulting model fits to a second model optimization analysis that followed the more traditional approach of fitting to summaries of accuracies across trial blocks (Love et al., 2004). In this approach, the average accuracy for both participant and model is calculated for each block of 16 trials and these block accuracies are used to calculate model fit error. In minimizing block-wise error between model and participant behaviour, this type of model optimization can account for patterns in learning accuracy across blocks (i.e., participant- and task-specific learning curves). We performed this second model optimization for each participant in each task and then used the optimized parameters to calculate the same log likelihood measure based on trial-by-trial responses as was used in the central model optimization. The logic follows that if trial-wise optimization provides a better account of behaviour (i.e., the specific responses made by participants on each trial), the trial-wise likelihoods will be significantly higher than the likelihoods from the block-wise optimization. Indeed, this is exactly what we found in comparing model fits between approaches in a mixed-effects regression: trial-wise optimization far exceeded block-wise optimization (mean *lnL_trial_*=-49.7, mean *lnL_block_*=-63.5, *t*(174)=4.56, *p*=9.7x10^-6^). These results suggest the trial-wise optimization provided a better account of participants’ specific responses on each trial and validates using the optimized parameters to generate trial-specific memory formation function predictions.

Best fitting parameters from the trial-wise optimization (mean and 95% confidence intervals: *γ* = 8.567 ± 1.374, β = 3.494 ± 0.327, η = 0.209 ± 0.049, *d =* 21.542 ± 3.532, *τ* = 0.191 ± 0.028) for each participant and task were then leveraged to generate model-based predictions for trials when memory clusters were created versus updated (**Figure 2A**). Trials were labelled as create or update trials according to the latent trial-wise operations of the model until the last cluster for a given task was created. At this point, memory modification was considered completed and the remaining trials were labeled as baseline. These predictions of create, update, and baseline trials served as model-based regressors for memory modification events in the fMRI analyses described below.

SUSTAIN predictions for memory operations during the learning tasks were consistent with prior reports (Love et al., 2004): The modal solution included 6 clusters (mean 6.17, standard deviation 1.5) with a minimum of 2 and maximum of 11 clusters (**Figure 2B**). Across all tasks, there was an average of 22.1 total create trials (range of 13-35 trials, 0.85 SE) and 36.4 total update memory trials (range of 23-67 trials, 2.25 SE) per participant.

### MRI data acquisition

Whole-brain imaging data were acquired on a 3.0T Siemens Skyra system at the University of Texas at Austin Imaging Research Center. A high-resolution T1-weighted MPRAGE structural volume (TR = 1.9s, TE = 2.43ms, flip angle = 9°, FOV = 256mm, matrix = 256x256, voxel dimensions = 1mm isotropic) was acquired for coregistration and parcellation. Two oblique coronal T2-weighted structural images were acquired perpendicular to the main axis of the hippocampus (TR = 13,150ms, TE = 82ms, matrix = 384x384, 0.4x0.4mm in-plane resolution, 1.5mm thru-plane resolution, 60 slices, no gap). High-resolution functional images were acquired using a T2*-weighted multiband accelerated EPI pulse sequence (TR = 2s, TE = 31ms, flip angle = 73°, FOV = 220mm, matrix = 128x128, slice thickness = 1.7mm, number of slices = 72, multiband factor = 3) allowing for whole brain coverage with 1.7mm isotropic voxels.

### MRI data preprocessing

Anatomical and functional volumes were preprocessed with fmriprep version 1.1.1 using default pipelines. We provide a brief summary of this pipeline here; the full methods of the pipeline can be found at http://osf.io/jk79v). Participant T1 volumes were skull-stripped and registered to the MNI 2009c asymmetric template using *ANTs* (Avants et al., 2011). Functional volumes were motion corrected (FSL *mcflirt*), corrected for susceptibility distortions using fieldmaps (FSL *fugue*), and registered to the participant’s T1 volume (Freesurfer *bbregister*). Functional volumes were resampled to MNI template space before the main analysis. Whole brain automated parcellation on each participant’s T1 volume was also conducted with Freesurfer (*recon-all*).

### Beta series estimation

For each participant, trial-level beta series were estimated for the first fMRI run in each learning task using the LS-S approach (Mumford et al., 2012b). Beta volumes for both stimulus presentation and feedback events were separately modelled for each trial; however, BOLD activity related to stimulus presentation was the focus of the current study. Confound regressors in the trial-level beta estimation included framewise displacement, six motion parameters (three degrees of translation and rotation) and estimates of BOLD signal noise as defined by the first six components of anatomical CompCor (Behzadi et al., 2007).

### fMRI analysis of memory formation events

To characterize neural signatures of memory formation events, whole-brain beta series of trial-by-trial stimulus presentations was analyzed with a mixed effects linear regression approach (*statsmodels* python library, version 0.8). All participants’ beta series from each task were concatenated and analyzed simultaneously with a custom searchlight kernel implemented in PyMVPA. Specifically, within each searchlight (radius=2 voxels), beta estimates were averaged and evaluated with a regression model that included trial-level fixed factors for: 1) model-predicted memory cluster modification events (create, update, baseline), 2) correct or incorrect responses, and 3) trial number. Participant was also included in the regression as a random intercept. Thus, the regression model was constructed to characterize neural activation that differed between model-predicted memory cluster create and update events relative to baseline while controlling for potentially confounding effects of response correctness and learning time. Applying this custom searchlight to the whole brain resulted in a statistical *t*-map of the contrast between create and update trials.

Cluster-level inferences for the create versus update contrast was performed by first estimating the noise model of the dataset and statistical analysis. The residual volumes from the regression analysis described above were extracted and analyzed with AFNI (Cox, 1996) *3dFWMHx* using the *acf* option to estimate the intrinsic autocorrelation of the data (*a*=0.659, *b*=3.082, *c*=11.479). These *acf* parameters were then inputted to AFNI *3dClustSim* (version 19.1.14) to estimate a cluster-extent threshold with 2-sided thresholding and third-nearest neighbour clustering. The result was a thresholding scheme in which the create vs. update statistical map was voxel-wise thresholded at *p*=0.001 and cluster corrected at *p*=0.05, which corresponded to a cluster-extent threshold of greater than 42 voxels. Regions of interest (ROIs) were defined from significant clusters and were labelled according to their spatial location relative to the Harvard-Oxford Structural or Juelich Histological atlases as included in FSL.

### Relating neural signatures of memory formation to learning

Time series were extracted from the trial-by-trial beta estimates for the hippocampal and ventral striatum ROIs, as well as any other significant cluster showing a create vs. update memory effect (**Table 2**). Specifically, the betas within a sphere defined by a radius of 2mm and centred on the voxel with the peak Z value for the contrast defining the cluster (see MNI coordinates in **Table 2**) were averaged for each trial and ROI. These time series were then used to calculate a neural index of the difference in create vs. update activation (i.e., memory modification effect) for each participant, task, and ROI.

**Table 2:** Significant clusters showing distinct neural engagement to model-derived memory formation events (i.e., memory create vs. update). All reported clusters showed greater activation for create relative to update events. Cluster information includes the corresponding anatomical label as defined in the Harvard-Oxford Structural or Juelich Histological atlases, the *t* statistic of the peak voxel, the cluster size in number of voxels, and peak voxel location in MNI coordinates (x, y, z).

| Region | Z peak | voxels | X | y | z |
| --- | --- | --- | --- | --- | --- |
| precuneus | 5.08 | 399 | -14 | -70 | 34 |
| right ventral striatum | 5.01 | 225 | 20 | 8 | -4 |
| left ventral striatum | 4.99 | 188 | -20 | 14 | -8 |
| premotor | 5.33 | 97 | -2 | 0 | 34 |
| superior parietal | 4.09 | 65 | -10 | -34 | 46 |
| right hippocampus | 5.26 | 57 | 30 | -18 | -19 |
| posterior cingulate | 4.18 | 54 | 6 | -38 | 26 |
| premotor | 4.71 | 44 | 6 | -14 | 48 |

We evaluated the relationship between the neural indices of the memory modification effect and end of learning performance with two complementary analyses. First, a mixed effects linear regression was conducted that included end of learning performance (tasks labeled as learners versus not), ROI, and their interaction as predictors of the memory modification effect with participants included as a random effect. Follow-up analyses that estimated the relationship between end of learning performance and create-update activation for each ROI in separate regression models were also conducted to aid interpretation.

We also performed a mixed effects binomial logistic regression analysis with the memory modification effect and ROI (specifically hippocampus and striatum regions) as predictors of end of learning performance (learner versus not) for each task with participants as a random effect (**Figure 3B**). With the binary outcome of task learner, this analysis is akin to a classification approach. Follow-up ROI-specific analyses were also conducted with mixed effects binomial logistic regression. The same ROI-specific analyses were also conducted for the premotor, precuneus, posterior cingulate, and superior parietal clusters.

**Figure 3.**
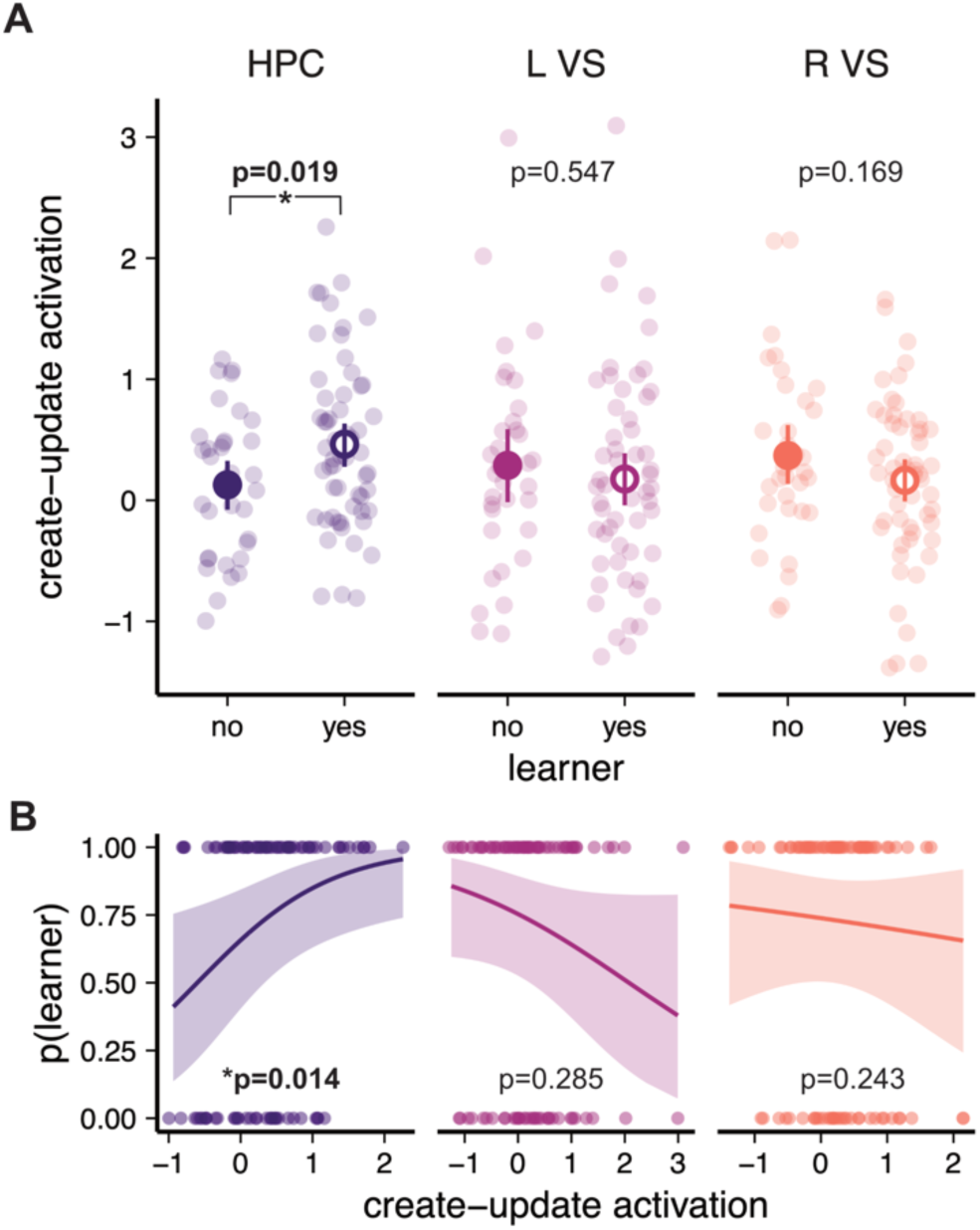
Neural indices of the memory computations that lead to successful learning outcomes. **A)** Activation differences that track memory creation and updating are plotted separately for tasks that reached learning criterion and those that did not (learner yes versus no) for hippocampus (HPC) and ventral striatum (VS). Group averages (larger circles) and bootstrapped 95% confidence intervals (dark lines) are depicted along with individual points for each completed task (lighter circles). **B)** Results of a logistic regression analysis assessing the relationship between whether activation reflected memory creation or updating and the likelihood that a task was successfully learned (“p(learner)”) are depicted with estimated marginal means of the statistical model (dark lines and confidence bands) and the individual task data (lighter circles). Data includes N=25 participants each in 4 tasks.

### Reliability of temporal coupling between model and neural measures

A key aspect of our approach is linking trial-wise model predictions of memory formation functions, as derived through simulations of learning behaviour, to neural measures. If task-specific model predictions of memory functions are capturing important dynamics in neural engagement, the observed differences between memory create and update trials should be significantly stronger than when the temporal coupling between model and brain is broken. To directly evaluated this, we conducted a permutation analysis in which model-based memory formation predictions were randomly shuffled across tasks within each participant and the primary analysis comparing neural engagement to memory create and update trials was performed. This procedure was repeated 1000 times to generate a distribution of estimated create-update trial effects (i.e., *z* statistics) separately for the HPC and striatum ROIs. A *p* value was calculated that corresponded to the proportion of shuffled analyses that resulted in a larger create-update effect than the observed effect.

### Controlling for stimulus and feature novelty

Throughout the learning tasks, participants were exposed to novel features and novel objects (i.e., novel combinations of features). Although this novelty is one factor in SUSTAIN’s mechanism for mismatching signaling, whether or not a memory is created or updated depends also on the learning experience and response history up to that point. Given the well-established effect of novelty signaling on memory function, especially in the hippocampus (e.g., Duncan et al., 2012), any effects observed in the current study may be driven more by this general novelty signaling rather than the learning-mediated mismatch signaling as formalized in SUSTAIN. As such, we characterized both model predictions of memory formation and neural activation to these memory operations during stimulus and feature novelty. Specifically, the same mixed effects linear regression model in the central analysis was conducted separately on trials that only included novel stimuli and trials that only included novel features. The logic of this control analysis follows that if SUSTAIN’s predictions of memory formation operations occur beyond that of simple novelty signaling, neural activation differences for create and update operations should persist during stimulus and feature novelty.

### Functional connectivity

To characterize potential functional networks associated with neural regions demonstrating distinct activation profiles during memory create versus update events, we conducted a functional connectivity analysis (Rissman et al., 2004). We targeted regions that showed learning-mediated activation as indexed by both a create vs. update memory effect and a link to learning outcome. This approach constrains the analysis such that learning behaviour and the corresponding model simulations act as a filter on relevant neural activation that we can then further characterize through functional connectivity. To conduct this analysis, the timeseries from the peak of a learning-related seed brain region was entered into a searchlight-based mixed effects linear regression analysis such that trial-by-trial activation interacted with the trial-level create versus update regressor to predict whole-brain trial-by-trial beta estimates. The regression model also included confound regressors for trial correctness and trial number. To additionally account for non-task related temporal noise in the beta series, we also included nuisance regressor timeseries from voxels within the ventricles (left: -18, -38, 15; right: 21, -38, 15) and white matter (left: -24, -5, 35; right: 24, -5, 35). A searchlight (radius=2 voxels) regression analysis was conducted on the whole brain to generate statistical *t*-maps that highlighted brain regions that were distinctly functionally coactive with the target seed region for memory create versus update trials. Significant clusters were identified in the same manner as the central analysis described previously with a cluster forming threshold of p=0.001 and cluster extent threshold of p=0.05, *k*=42 voxels.

## Results

### Neural indices of learning-guided memory formation

Participants learned to varying degrees across the four tasks (**Figure 1D**) with end of learning performance splitting roughly equally into tasks in which learning was successful (>90% accuracy, 57/100 tasks) or not (<90% accuracy, 43/100 tasks). To characterize how different memory operations—creation and updating—guided successful learning performance, model-based predictions of when distinct memory functions occurred throughout early stages of learning were derived from participant and task-specific simulations of SUSTAIN. For each trial, the model output a label indicating whether a new memory was created or an existing knowledge cluster updated (**Figure 2A**). We then identified brain regions distinctly associated with each memory mechanism by conducting a voxel-wise mixed-effects regression on whole-brain trial-by-trial betaseries (Mumford et al., 2012a). Specifically, at each voxel we estimated how computational model-derived memory create versus update trials were associated with changes in neural activation.

Notably, an anterior region of the right hippocampus (HPC, peak MNI coordinate [30, - 18, -19], 57 voxels, voxel-wise threshold p = 0.001, cluster-extent threshold p = 0.05) and bilateral regions of ventral striatum (left: L VS, peak [-20, 14, -8], 188 voxels; right: R VS, peak [8, 20, -4], 225 voxels) demonstrated distinct neural engagement for create versus update trials (**Figure 2C/D**); all three regions showed more engagement when new knowledge clusters were created relative to when clusters were updated. Importantly, the effects in both anterior hippocampus (p = 0.005) and right ventral striatum (p = 0.022), but not left ventral striatum (p = 0.232), were robust to within-participant random permutation tests (**Figure 2C/D**, violin plot insets). Additional regions across the brain similarly tracked when new memories were created and existing memories updated, including precuneus, premotor cortex, and superior parietal cortex (**Table 2**). Importantly, these neural signatures of distinct memory operations were independent of trial outcome or number and were evident across all tasks, each with their own stimulus sets; thus, ruling out explanations based on error-related and stimulus-specific effects. Also, separate control analyses restricted to trials with novel stimuli or novel features showed significant activation differences between cluster creation and updating in all three ROIs (all p<0.01), suggesting that the observed neural effects are due to processes beyond detecting novel features or novel combinations of features. Rather, these findings demonstrate that fundamentally different memory operations, related to the formation of category structures and as formalized in computational predictions of SUSTAIN, are associated with unique neural signatures in key learning regions like hippocampus and striatum.

### Hippocampal engagement to memory operations predicts accurate categorization decisions

Given their theorized role in learning (Antzoulatos and Miller, 2011; Davis et al., 2012b; Ballard et al., 2018; Mack et al., 2018; Calderon et al., 2021), we next assessed the degree that the distinct memory operations supported by anterior hippocampus and ventral striatum were associated with categorization accuracy. We reasoned that the more a participant showed distinct engagement to create versus update trials early in learning, the better their learning outcome. In line with this prediction, a linear mixed effects regression across the ROIs resulted in a main effect of learner (*χ*^2^=4.340, p=0.037), but also an interaction with ROI (*χ*^2^=6.457, p=0.039). Follow-up analyses separated by ROI showed that for tasks in which participants were highly accurate by the end of learning (>90% accuracy), anterior hippocampus activation during early learning exhibited a larger difference in memory create versus update trials (**Figure 3A**; β=0.336, CI=[0.056, 0.616], p=0.019). No such effect was found in either ventral striatum cluster (L VS: β=-0.113, CI=[-0.479, 0.254], p=0.547; R VS: β=-0.206, CI=[-0.499, 0.088], p=0.169). In a complementary analysis, we performed a logistic regression to predict whether the create versus update activation difference early in learning predicted accurate categorization decisions at the end of learning (**Figure 3B**). Applying this analysis to all ROIs showed a main effect of memory modification (χ^2^=5.726, p=0.017) and an interaction with ROI (χ^2^=8.276, p=0.016). Follow-up ROI-specific regressions revealed that these effects where driven by anterior hippocampus activation predicting learning outcome (log odds=2.752, CI=[1.225, 6.181], p=0.014), while ventral striatum did not (L VS: log odds=0.742, CI=[0.429, 1.282], p=0.285; R VS: log odds=0.663, CI=[0.333-1.322], p=0.243). Additionally, we performed the same learner vs. non-learner and logistic regression analyses for the additional clusters in premotor, precuneus, posterior cingulate, and superior parietal cortex (see **Table 2**); none of these regions showed a significant relationship between the create vs. update memory effect and learning outcome (all ps>0.08). These findings point to a specific role for hippocampus in mediating when individuals exploit existing knowledge and when they create new knowledge. The more distinctly anterior hippocampus was engaged for memory create versus update events in the initial stages of learning, as predicted by SUSTAIN, the more likely that participants reached high categorization accuracy at the end of learning.

### Networks supporting memory creation and updating

With evidence that anterior hippocampus activation is linked to qualitatively different memory operations and that the degree of this engagement distinction leads to successful learning outcomes, we reasoned that distinct cortical networks may work in concert with anterior hippocampus to support memory creation versus updating early in learning. Indeed, memory creation is triggered by surprising events and results in a detailed encoding of the current experience into conceptual knowledge that influences subsequent category decisions. As such, hippocampal memory create mechanisms may be supported by frontoparietal regions, including lateral parietal cortex areas sensitive to novelty (Uncapher and Wagner, 2009; Hutchinson et al., 2014) and detailed memory encoding (Kuhl and Chun, 2014) and medial prefrontal cortex (mPFC) regions that encode goal-relevant memory models (van Kesteren et al., 2010; Preston and Eichenbaum, 2013; Schlichting and Preston, 2015, 2016; Mack et al., 2016, 2020; Gilboa and Marlatte, 2017; Bowman and Zeithamova, 2018; Theves et al., 2021).

To investigate these possibilities, we performed an interregional functional correlation analysis (Rissman et al., 2004) to identify brain regions that were distinctly coactive with anterior hippocampus during memory create versus update trials. A linear mixed effects regression analysis that related the functional time series of the right anterior hippocampus cluster to the rest of the brain as a function of which memory computation was predicted by SUSTAIN on a given trial revealed two distinct sets of regions (**Figure 4**). Subgenual vmPFC (peak Z=4.31, [-8, 24, -10], 78 voxels) and right angular gyrus (peak Z=4.1, [48, -42, 22], 47 voxels) were significantly more correlated with anterior hippocampus during memory create versus update trials. In contrast, three regions in close proximity within premotor cortex (peak Z=4.67, [-16, -22, 76], 161 voxels; peak Z=4.21, [12 -18, 70], 160 voxels; peak Z=3.83, [-12, -8, 78], 57 voxels) exhibited greater correlation with anterior hippocampus during memory update versus create trials. Importantly, these interregional correlation effects were observed independent of response accuracy and trial number. Moreover, none of these regions showed distinct overall engagement to memory formation functions. Thus, that trial-by-trial neural dynamics in anterior hippocampus are distinctly reflected in these brain regions during specific memory formation events supports the proposal that hippocampal learning occurs in coordination with key frontoparietal networks (Xue et al., 2013; Hutchinson et al., 2014; Braunlich et al., 2015; Schlichting and Preston, 2016; Bowman and Zeithamova, 2018; Morton et al., 2020; Ashby and Zeithamova, 2022).

**Figure 4.**
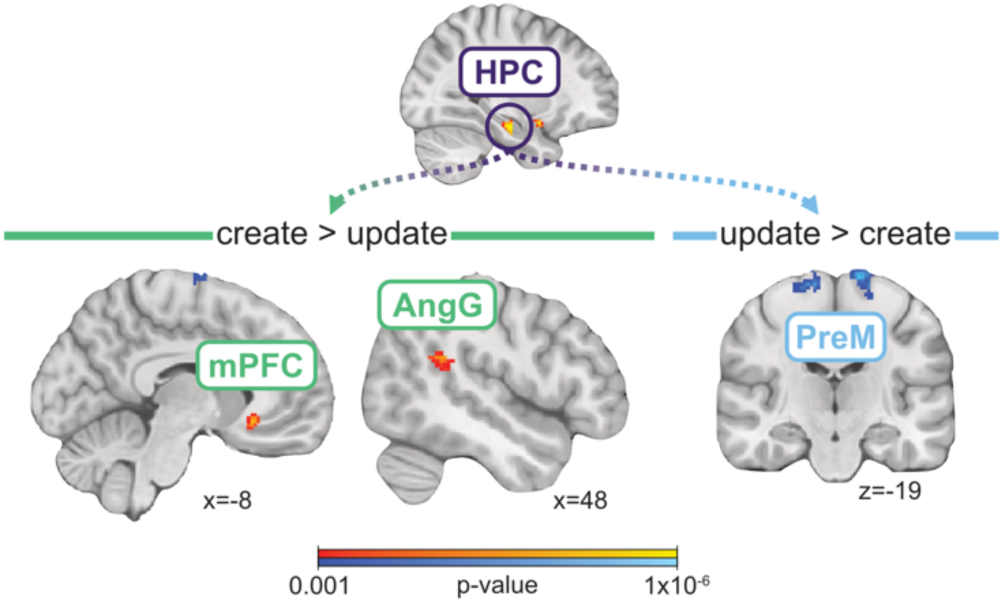
Interregional functional correlation with HPC. Regions of mPFC and angular gyrus showed greater functional coactivation with HPC during memory create relative to update trials. Regions of premotor cortex exhibited greater correlation with HPC during update relative to create trials. Clusters were defined by a voxel wise threshold of p=0.001 and cluster-extent threshold of p=0.05, which corresponded with a cluster extent of 42 voxels. N=25.

## Discussion

Our neurocomputational approach isolates moments of qualitatively distinct memory operations that lead to successful learning outcomes. We show that when anterior hippocampus is engaged during moments of new concept creation, as opposed to updating existing concepts, participants succeed at learning. In contrast, ventral striatum engagement tracks the shift between memory creation and updating but does not predict learning outcomes. These results suggest that anterior hippocampus, together with vmPFC and angular gyrus, forms distinct knowledge clusters that differentiate concepts, supporting rapid acquisition of category knowledge and accurate decision making (Schapiro et al., 2017; Mack et al., 2018; Sučević and Schapiro, 2023).

Classical views posit distinct memory systems involved in concept learning: a hippocampal-based declarative system that encodes initial episodic memories of concept exemplars, and a basal ganglia-based procedural system that underlies implicit stimulus-response (i.e., category labels) learning (Knowlton et al., 1996; Squire and Zola, 1996; Shohamy et al., 2008). Our findings refine this view. Although ventral striatum tracked model-predicted shifts in memory operations, it did not relate to individual differences in learning. The more rigid striatal learning system may be ill-equipped for learning the complex multidimensional associations required in the current tasks. In contrast, the hippocampus provides a flexible learning system capable of building multidimensional category representations (Davis et al., 2014; Mack et al., 2016; Bowman and Zeithamova, 2018; Sučević and Schapiro, 2023). Indeed, that hippocampal activation during early create and update trials predicted learning outcome offers compelling evidence for such an account.

By characterizing learning with model-based predictions of memory operations, our findings significantly extend recent findings establishing hippocampus as central to concept learning. It has been demonstrated in humans (Mack et al., 2016; Bowman and Zeithamova, 2018; Bowman et al., 2020; Theves et al., 2020) and models (Heffernan et al., 2021; Sučević and Schapiro, 2023) that hippocampal activation patterns reflect the latent structure of multidimensional category spaces, yet these results largely depend on neural coding at the end of learning. Also, measures of category evidence throughout learning have been linked to trial-by-trial hippocampal activation (Davis et al., 2012a; Theves et al., 2021), but the mechanisms driving these effects have remained unclear. Here, we establish that anterior hippocampal engagement during new concept learning is mediated by shifts between qualitatively different memory create and update operations.

More broadly, our findings align with evidence of novelty signaling (Kumaran and Maguire, 2009; Duncan et al., 2012; Larkin et al., 2014; Bein et al., 2020; Sinclair et al., 2021) and rapid remapping in hippocampus during spatial navigation (Steemers et al., 2016; Julian and Doeller, 2021; Zheng et al., 2021), episodic memory (Sinclair et al., 2021), associative memory (Wanjia et al., 2021), and event segmentation (Baldassano et al., 2017; Clewett et al., 2019). Our results provide unique support for a similar rapid encoding mechanism during category learning, in which distinct hippocampal states reflect different memory operations (Duncan et al., 2012; Bein et al., 2020). Coordination of these states is key to building adaptive knowledge structures that capture regularities and discriminate novel experiences to enhance learning outcomes (Love et al., 2004; Mack et al., 2018; Sučević and Schapiro, 2023).

Importantly, our findings were possible only by linking learning behaviour to neural activation with a computational model. Leveraging SUSTAIN to simulate participant-specific behaviour allowed us to identify key shifts between the model’s memory creation and updating functions. Notably, this participant- and task-specific mapping between behaviour, model predictions, and neural measures was essential—the create greater than update effect in hippocampus was significantly stronger than permutation analyses that shuffled task mappings within participants. Two additional aspects of our study strengthen the results: a) the memory modification effect generalized across four learning tasks, arguing against stimulus-specific factors; and b) most participants successfully learned some but not all tasks (19 mixed, 5 all, and 1 none), allowing for a stronger within-participant test of neural signatures linked to learning outcome.

That hippocampal engagement during learning reflected the dynamics of memory operations in SUSTAIN (Love et al., 2004) affords a direct mechanistic interpretation. SUSTAIN proposes that the degree of match between current experience and stored knowledge dictates the learner’s response. But unlike other successful categorization models that rely on fixed representational formats (Homa et al., 1973; Nosofsky, 1986; Smith and Minda, 1998), SUSTAIN uniquely posits that this match also drives how stored knowledge is modified—a high match signals an update to an existing memory trace, whereas a low match triggers creation of a new trace for the current experience. Learning to discriminate a categorization structure requires an optimal number of clusters that generalize to new experiences to varying degrees. SUSTAIN’s memory create and update operations, coupled with selective attention to stimulus dimensions, build concept knowledge that reflects regularities while also supporting accurate discrimination for the learning goal. The current findings are consistent with and extend mounting evidence linking SUSTAIN’s formal mechanisms to neural measures in humans (Love and Gureckis, 2007; Davis et al., 2012a; Mack et al., 2016, 2018, 2020; Braunlich and Love, 2019) and animals (Broschard et al., 2021b, 2021a).

Hippocampal interactions with cortical structures during critical learning moments are also reflected in our findings. Hippocampal-prefrontal interactions are known to support the formation of structured memory networks (Gilboa and Marlatte, 2017; Morton et al., 2017) that underlie complex associative learning (Schlichting et al., 2015; Schlichting and Preston, 2016), schemas (Zheng et al., 2021), and categories (Constantinescu et al., 2016; Mack et al., 2016; Bowman and Zeithamova, 2018; Theves et al., 2021). Accordingly, we predicted and observed that during initial learning, experiences warranting the creation of a new memory trace showed increased hippocampus-PFC coactivation. This supports the view that hippocampal mismatch signals may influence schema abstraction in mPFC (Robin and Moscovitch, 2017) to highlight information most diagnostic for current task goals (Bowman and Zeithamova, 2018; Mack et al., 2020).

We also found stronger hippocampal coactivation with ventral posterior parietal cortex during memory creation. While parietal cortex is well known for its role in memory retrieval (Wagner et al., 2005; Hutchinson et al., 2014; Kuhl and Chun, 2014; Morton et al., 2020; Zhao et al., 2021), ventral posterior regions also play an important role during encoding. As part of the ventral attention network, these regions reorient attention to unexpected but behaviourally relevant information during encoding (Corbetta et al., 2008; Uncapher and Wagner, 2009; Turk-Browne et al., 2013). Moreover, angular gyrus engagement supports integration of event elements and vivid subsequent memory (Tibon et al., 2019). These encoding-based functions likely supported learning in the current paradigm— orienting to new feature combinations promotes rapid encoding of distinct memory traces. Our finding that ventral posterior parietal cortex tracks hippocampal engagement during memory creation provides unique evidence that these regions jointly support rapid learning of information that diverges from prior experience.

We additionally observed that premotor cortex areas were functionally more coactive with hippocampus during memory updating than creation. Premotor cortex is a key player in COVIS, a prominent multiple learning systems account (Ashby et al., 1998; Ashby and Maddox, 2005). Specifically, an implicit procedural learning circuit between striatum and premotor cortex is theorized to encode direct stimulus-response associations. That we found coactivation between premotor cortex and hippocampus rather than striatum runs counter to these theories. However, hippocampal place coding (O’Keefe and Nadel, 1978; Eichenbaum et al., 1999; Foster and Knierim, 2012), in which behaviourally relevant spatial sequences are stored and activated to support action plans, suggests that hippocampus encodes motor alongside sensory information. The hippocampus-premotor coactivation we observed may reflect strengthening of associations between the current stimulus and motor response, a situation likely to occur when existing memories are updated. These results motivate further inquiry into how multiple learning systems coordinate during early learning.

One limitation to the current paradigm is that each category structure was defined by only eight unique stimuli (Shepard et al., 1961). This may have encouraged explicit learning processes, which may be biased towards episodic-like functions of hippocampus (Squire and Zola, 1996). Conversely, category structures with many exemplars are thought to engage implicit learning mechanisms tied to striatal function (Ashby and Maddox, 2005; Minda et al., 2024). Future work investigating category and exemplar structure, and implicit versus explicit learning would further clarify how multiple systems drive different learning experiences.

In summary, our findings propose a key revision to long-held theories of concept learning (Knowlton et al., 1996; Squire and Zola, 1996; Shohamy et al., 2008) such that hippocampus is an important player in building organized memories from learning’s earliest moments. Although we found unique signatures of complementary memory formation functions during early learning in hippocampus and striatum, only the hippocampal signature of memory modification was associated with learning outcome. These results are consistent with neurobiological theories of hippocampus (Rolls, 2013; Schapiro et al., 2017; Sučević and Schapiro, 2023) and formal cognitive models (Love et al., 2004), and provide novel support for the theoretical convergence of these two perspectives (Love and Gureckis, 2007; Mack et al., 2018). By leveraging trial-specific model predictions of latent memory operations, we identified theoretically meaningful learning moments and characterized the neural mechanisms that support the formation of flexible concept knowledge.

## Acknowledgements

The authors thank Meg Schlichting, Katherine Duncan, and Dasa Zeithamova for helpful discussions in preparation of this manuscript. This work was supported by the National Institute of Mental Health (F32-MH100904 to MLM, R01-MH100121 to ARP), Natural Sciences and Engineering Research Council (Discovery Grants RGPIN-2017-06753 and RGPIN-2024-0588 to MLM), Canada Foundation for Innovation and Ontario Research Fund (36601 to MLM), Economic and Social Research Council (ES/W007347/1 to BCL), and a Royal Society Wolfson Fellowship (18302 to BCL).

